# Innervation of Retrodiscal Tissues in Patients with Temporomandibular Joint Disorder

**DOI:** 10.1101/2025.06.11.659119

**Authors:** Jessie J. Alfaro, Anahit H. Hovhannisyan, Erin E. Locke, RE-JOIN Consortium Investigators, Felix J. Amarista, Daniel E. Perez, Armen N. Akopian

## Abstract

Temporomandibular joint (TMJ) disorders (TMJDs) are a group of musculoskeletal conditions affecting the orofacial region and often associated with facial pain. Understanding the sensory innervation of TMJ structures, particularly the retrodiscal tissue, is essential for identifying pain mechanisms in TMJD because to study these mechanisms, we must first determine the sensory neuronal makeup of the TMJ. However, data on nerve types within TMJ tissues remain limited. This study examined the sensory and sympathetic nerve profiles in retrodiscal tissues from TMJD patients with osteoarthritis (OA), rheumatoid arthritis (RA), or condylar hyperplasia (CH) who underwent bilateral TMJ replacement (TMJR). Immunohistochemistry with specific nerve markers was used to visualize and quantify nerve subtypes. CH tissues had significantly lower densities of pgp9.5^+^ sensory fibers compared to RA and OA, which showed similar levels. Across all subtypes, the ratio of unmyelinated (pgp9.5^+^/NFH^−^) to myelinated (pgp9.5^+^/NFH^+^) fibers was approximately 70:30. Most sensory nerves were CGRP^+^ (peptidergic), while a smaller portion were CGRP^−^ (non-peptidergic), some of which were parvalbumin-positive (PV^+^). Both myelinated and non-myelinated peptidergic as well as non-peptidergic fibers were present in the retrodiscal tissues. In addition to sensory innervation, all retrodiscal tissues contained tyrosine hydroxylase-positive (TH^+^) sympathetic fibers, primarily innervating blood vessels (alpha-smooth muscle actin (α-SMA^+^)). These vessels were also predominantly innervated by unmyelinated sensory fibers, with limited input from myelinated sensory nerves. In summary, all TMJD subtypes shared similar nerve compositions, but CH tissues exhibited reduced sensory nerve density, a potential explanation for the lower association with pain compared to OA and RA. For all TMJD subtypes, retrodiscal tissue vasculature was mainly innervated by sympathetic and unmyelinated sensory nerves. These findings enhance understanding of the neural basis of TMJD-related pain.

## Introduction

Temporomandibular joint (TMJ) disorders (TMJDs) represent a group of joint and musculoskeletal conditions affecting approximately 30 million individuals in the United States alone (Romero-Reyes and Uyanik 2014; Tmd (temporomandibular disorders) 2025). Among those affected with TMJDs, pain localized to the TMJ is reported by approximately 40% of patients, with about 15% experiencing chronic pain (Facial pain 2018; Prevalence of tmjd and its signs and symptoms). Notably, TMJD-associated pain can be alleviated through local anesthetic blockade, highlighting the role of peripheral sensory innervation in TMJD pathophysiology (Danzig et al. 1992). Considering the relevance of TMJ sensory neurons to TMJDs, a comprehensive understanding of TMJ sensory innervation is essential for elucidating pain mechanisms in TMJD.

Most existing studies on TMJ innervation have focused on the synovial lining layer, a component of the joint capsule. Early investigations into TMJ synovial innervation were conducted in animal models, particularly mice and non-human primates (Frommer and Monroe 1966; Keller and Moffett 1968). These studies identified peptidergic sensory nerves within the TMJ (Ichikawa et al. 1989; Johansson et al. 1986; Kido et al. 1993). Further electrophysiological studies revealed that the TMJ is innervated by two primary nociceptive afferent subtypes: Aδ-fibers and C-fibers (Takeuchi et al. 2001; Takeuchi and Toda 2003).

In humans, data on TMJ innervation have been derived from both anatomical and electrophysiological investigations. The primary source of TMJ sensory innervation is the auriculotemporal nerve (ATN), a branch of the mandibular division (V3) of the trigeminal nerve (*Fig 1*) (Greenberg and Breiner 2025; Kucukguven et al. 2022; Schmidt et al. 1998). Using silver staining techniques, both myelinated and unmyelinated nerve fibers have been visualized in the anterior portion of the TMJ disc (Asaki et al. 2006). Microneurography recordings from single ATN fibers in adult human subjects identified both slow- and fast-adapting sensory units, encompassing Aδ and C fibers (Ishikawa 1989).

**Figure 1.**
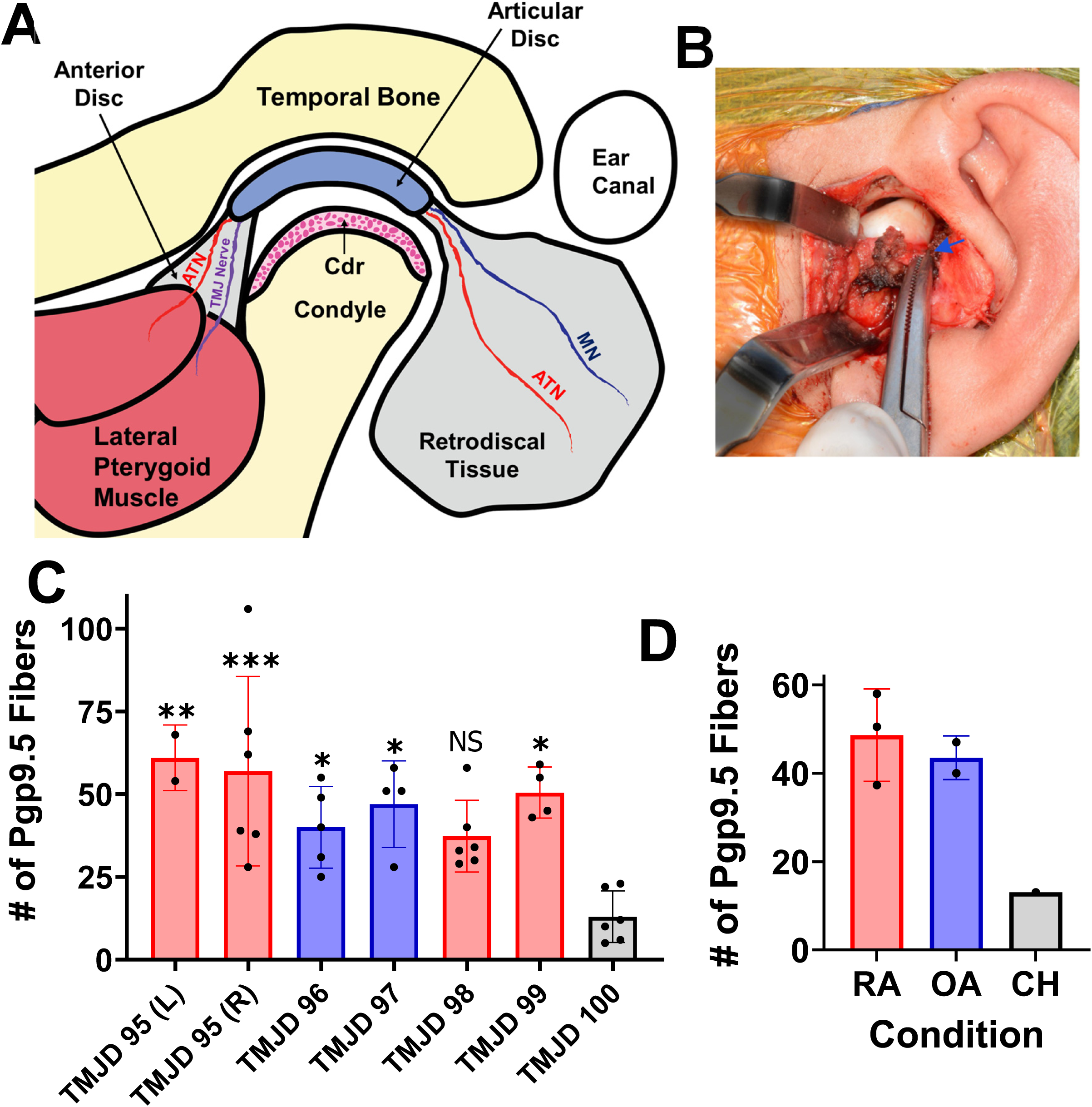
Sensory innervation of retrodiscal tissue in TMJD subtypes. (**A**) Schematic representation of the temporomandibular joint (TMJ) and its sensory innervation. The TMJ is innervated by the auriculotemporal nerve (ATN), masseteric nerve (MN), and other TMJ-specific nerve branches. The chondrocyte layer of the mandibular condyle is labeled as “Cdr”. (**B**) Intraoperative image from total joint replacement (TMJR) in a TMJD participant. The retrodiscal tissue, excised for analysis, is indicated by the blue arrow. (**C**) Quantification of pgp9.5^+^ sensory nerve fibers in retrodiscal tissue from individual TMJD participants. Bars for participant #95 represent data from left (L) or right (R) retrodiscal tissue sections. Participants with rheumatoid arthritis (RA) are shown in red, osteoarthritis (OA) in blue, and condylar hyperplasia (CH) in grey. N represents the number of cryosections analyzed per tissue. Statistical analysis was performed using one-way ANOVA with Bonferroni’s post hoc test (*p < 0.05; **p < 0.01; ***p < 0.001). (**D**) Grouped analysis of pgp9.5^+^ sensory fiber density across TMJD subtypes. N indicates the number of participants per group.

Despite these insights, there remains a paucity of detailed information regarding the innervation of specialized human TMJ structures, including the capsular tissue with its synovial lining, the articular disc, anterior disc, and retrodiscal tissue. The current study aimed to address this gap in knowledge by characterizing the types of sensory and sympathetic nerves present in the retrodiscal tissues obtained from patients with osteoarthritis, rheumatoid arthritis, or condyle hyperplasia TMJD who underwent bilateral TMJ replacement (TMJR).

## Materials and Methods

### Patient cohort and clinical assessment

This study included six female patients, aged 19 to 63 years, who were diagnosed with osteoarthritis, rheumatoid arthritis, or condyle hyperplasia TMJD. All patients underwent bilateral TMJR at the University Hospital in San Antonio, Texas. Retrodiscal tissue samples were collected intraoperatively during the TMJR procedures from participants enrolled in the “Retrodiscal, Capsular, and Discal TMJ Tissue Analysis and Repository” study (IRB #: HSC20220807H) between 2023 and 2025. Prior to surgery, participants underwent a comprehensive clinical evaluation comprising a battery of validated assessment tools. These included: Brief Pain Inventory – Short Form (BPI-SF); Tobacco, Alcohol, Prescription Medications, and Other Substances (TAPS); Pain–Sleep Duration Questionnaire; Pain, Enjoyment of Life, and General Activity Scale (PEG); Pain Catastrophizing Scale; Visual Analog Scale (VAS) for Pain; Douleur Neuropathique 4 (DN4) Questionnaire; Patient Health Questionnaire-2 (PHQ-2); Patient Global Impression of Change (PGIC); Patient-Reported Outcome Measurement Information System – Physical Function 6b (PROMIS SF v1.2); Patient-Reported Outcome Measurement Information System – Sleep Disturbance 6a (PROMIS SF v1.0), and Generalized Anxiety Disorder-2 (GAD-2). This comprehensive pre-operative assessment provided a multidimensional profile of each patient’s pain experience, psychosocial status, functional capacity, and overall well-being, thereby supporting the clinical context for tissue analysis.

### Retrodiscal tissue procurement and cryosectioning

Retrodiscal tissues were obtained from left and/or right TMJ. These retrodiscal tissue samples were fixed overnight in 4% paraformaldehyde (PFA) at 4°C. The following day, tissues were rinsed twice with 1× phosphate-buffer (PB) and subsequently cryoprotected in 10% sucrose for 18–24 hours, followed by 30% sucrose for a minimum of 18 hours at 4°C. After cryoprotection, tissues were embedded in Neg-50™ embedding medium (Fisher Scientific, Cat# 22-110-617) and sectioned at a thickness of 30–35 μm using a cryostat (TN55, Tanner Scientific).

### Immunohistochemistry (IHC) procedure

Immunostaining was performed as previously described (Hovhannisyan et al. 2023). Briefly, tissue sections were blocked for 90 minutes at room temperature (RT) in a solution containing 4% normal donkey serum (Sigma, St. Louis, MO), 2% bovine gamma-globulin (Sigma-Aldrich, St. Louis, MO), and 0.3% Triton X-100 (Fisher Scientific) in 0.1 M phosphate-buffered saline (PBS). Following blocking, sections were incubated overnight at RT with primary antibodies. After incubation, sections were washed with 0.1 M PBS to remove any unbound primary antibodies and then incubated for 90 minutes at RT with appropriate fluorophore-conjugated secondary antibodies (1:200 dilution; Jackson Immuno-Research, West Grove, PA, USA). Finally, slides were washed 3 × for 5 minutes with 0.1 M PBS, air-dried, and covered with Vectashield Antifade Mounting Medium (Vectorlabs SKU H-1000-10).

The following primary antibodies, previously validated on non-human primate tissues (Hovhannisyan et al. 2023; Tram et al. 2023), were used for human tissue sections: anti-neurofilament heavy chain (NFH) chicken polyclonal antibody (BioLegend, catalog #PCK-592P, 1:300); anti-PGP9.5 rabbit polyclonal antibody (Millipore-Sigma, #AB1761-I, 1:400); anti-CGRP guinea pig polyclonal antibody (Synaptic Systems, Goettingen, Germany, catalog #414 004, 1:200); anti-MrgprD rabbit polyclonal antibody (Alamone Labs, ASR-031, 1:200); anti-tyrosine hydroxylase (TH) rabbit polyclonal antibody (Pel-Freez, Rogers, AR, catalog #P40101, 1:400), and anti-smooth muscle actin (α-SMA) Cy3-conjugated mouse monoclonal antibody (Sigma, catalog #C6198, 1:200). Secondary antibodies were all raised in donkey (Jackson Immuno-Research).

### Evaluation of retrodiscal sections, imaging and fiber counting

Entire retrodiscal sections were visually assessed under a fluorescent microscope. The average number of PGP9.5^+^ sensory nerve fibers per field of view (FOV) was estimated. Several randomly selected FOV images were captured for each tissue, which generated sections for 20-50 slides (2 sections per slide). Images were acquired using a Keyence BZ-X810 all-in-one microscope (Keyence, Itasca, IL, USA) utilizing the Z-stack/“sectioning” function. Z-stack IHC images were captured for subsequent fiber counting. Imaging was performed with either a 10× or 20× objective. Control immunohistochemistry (IHC) was conducted on tissue sections processed as described, but with the omission of either primary antibodies or both primary and secondary antibodies. Imaging settings were optimized to ensure that the negative control conditions - lacking primary antibodies, and both primary and secondary antibodies - yield little-to-no possible positive nerve fiber signal.

Fiber counting was performed manually as previously described to quantify the total number of peripheral nerve types in FOV of retrodiscal tissues (Hovhannisyan et al. 2023). The rationale for using manual fiber counting to detect and quantify nerve fibers within FOV has been thoroughly discussed in earlier studies (Hovhannisyan et al. 2023; Tram et al. 2023).

### Statistical Analyses

Statistical analyses were performed using GraphPad Prism 10 (GraphPad Software, La Jolla, CA). Data are presented as mean ± standard deviation (SD). Correlations between nerve subtypes and pre-operative clinical examination data were not analyzed. Meaning of N – number is specified in the text for each dataset. Differences between participants or groups were assessed using unpaired t-tests or one-way analysis of variance (ANOVA) with Bonferroni’s post-hoc test. Statistical significance was accepted at p < 0.05. Interaction F ratios and the associated p-values are reported.

## Results

### Sensory innervation of retrodiscal tissue in TMJD subtypes

Retrodiscal tissues were collected using TJR procedures from female participants diagnosed with osteoarthritis (OA), rheumatoid arthritis (RA), or condylar hyperplasia (CH) TMJD. These collected samples were used for analysis of sensory innervation (*Figs. 1, 2*). Each retrodiscal specimen generated cryosections mounted on 20-50 slides, with 2-3 tissue sections per slide. A minimum of four randomly selected slides per tissue were immunolabeled with pgp9.5, a pan-sensory neuronal marker (Hovhannisyan et al. 2023).

**Figure 2.**
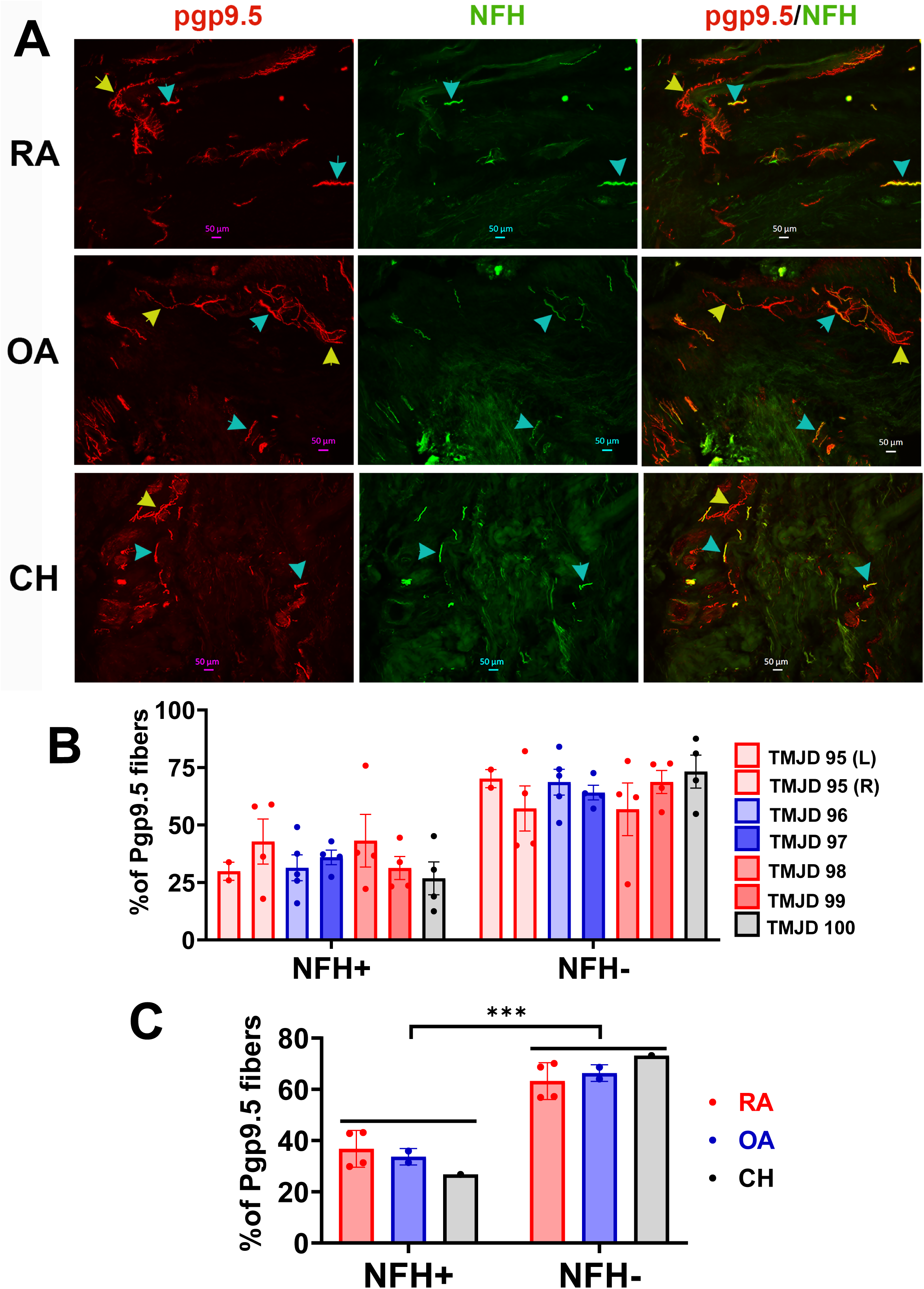
Myelinated and non-myelinated sensory fibers in retrodiscal tissue of TMJD subtypes. **(A)** Representative immunofluorescence micrographs showing NFH^+^ (myelinated A-fibers) and pgp9.5^+^ (pan-sensory neuronal marker) sensory nerve fibers in retrodiscal tissue sections from TMJD participants diagnosed with RA (top row), OA (middle row), and CH (bottom row). Cyan arrows indicate pgp9.5^+^/NFH^+^ fibers (myelinated), while yellow arrows indicate pgp9.5^+^/NFH^−^ fibers (unmyelinated). Antibodies and their corresponding fluorescence channels are noted in each image. Scale bars are included in each panel. (**B**) Quantification of pgp9.5^+^/NFH^+^ and pgp9.5^+^/NFH^−^ fibers in retrodiscal tissue from individual TMJD participants. Bars for participant #95 show counts from both left (L) and right (R) retrodiscal tissues. RA cases are shown in red, OA in blue, and CH in grey. N represents the number of frames analyzed per tissue. (**C**) Grouped analysis of pgp9.5^+^/NFH^+^ and pgp9.5^+^/NFH^−^ fiber percentages across TMJD subtypes. N indicates the number of tissues (or participants) analyzed per group. Statistical significance was determined using two-way ANOVA with Bonferroni’s post hoc test (***p < 0.001).

Entire slides were systematically examined using a “tile motion” scanning approach under a 10× objective, and pgp9.5-positive fibers were manually counted across complete tissue sections. The fiber counts were then averaged per field of view (FOV). Regions lacking pgp9.5^+^ fibers were recorded as zero counts. Variability in pgp9.5^+^ fiber density across retrodiscal tissues was minimal among OA and RA participants (*Fig. 1C*). However, the retrodiscal tissue from the participant with CH exhibited a significantly lower density of sensory fiber innervation (one-way ANOVA; F(6, 26) = 5.174; p = 0.0013; n = 2-6 sections per tissue; *Fig. 1C*). When data was grouped by TMJD subtype, retrodiscal tissues from OA and RA participants exhibited at least a threefold higher sensory fiber density (pgp9.5^+^) compared to the CH sample (RA: 48.60 ± 10.48; OA: 43.50 ± 4.95; CH: 13; *Fig. 1D*). Overall, retrodiscal tissues from TMJD participants demonstrated the presence of sensory nerve innervation, with significantly greater innervation density observed in OA and RA cases compared to CH.

### Myelinated and unmyelinated sensory fiber composition in retrodiscal tissues from TMJD subtypes

To further characterize the sensory innervation of retrodiscal tissue, we examined the ratio of myelinated to unmyelinated sensory nerve fibers across TMJD subtypes. Neurofilament heavy chain (NFH) was used as a marker of myelinated fibers. For each retrodiscal tissue sample, sections exhibiting the highest density of pgp9.5^+^ labeling were selected for analysis. Sensory fibers co-expressing pgp9.5 and NFH (pgp9.5^+^/NFH^+^) were classified as myelinated A-fibers, whereas fibers positive for pgp9.5 but lacking NFH (pgp9.5^+^/NFH^−^) were classified as unmyelinated C-fibers.

Representative images illustrating A-fibers (cyan arrows) and C-fibers (yellow arrows) in retrodiscal sections from RA, OA, and CH TMJD participants are shown in *Figure 2A*. Quantification of fiber subtypes revealed a consistent distribution across individual participants, with approximately 70% of fibers identified as C-fibers and 30% as A-fibers (*Fig. 2B*). When data were grouped by TMJD subtype, a similar trend was observed: retrodiscal tissues from RA, OA, and CH participants all exhibited a significantly higher proportion of C-fibers compared to A-fibers (one-way ANOVA; ***p < 0.001; *Fig. 2C*). Overall, although the absolute density of sensory fibers was lower in retrodiscal tissue from the CH participant, the ratio of C- to A-fibers remained consistent across all TMJD subtypes. Moreover, our findings indicate that unmyelinated C-fibers represent the predominant sensory fiber type innervating retrodiscal tissues from TMJD patients.

### Peptidergic sensory fibers in retrodiscal tissues from TMJD subtypes

Peptidergic sensory fibers were identified using immunohistochemistry (IHC) with CGRP antibodies, as previously described (Hovhannisyan et al. 2023). CGRP labeling was substantially weaker than that observed with pgp9.5 or NFH (*Fig. 3A* vs. *Fig. 2A*), particularly in retrodiscal tissue from CH TMJD participants (*Fig. 3A, bottom panels*). CGRP signal intensity varied among TMJD participants. In addition to sensory fibers, CGRP also labeled blood vessels and a subset of retrodiscal tissue cells (*Fig. 3A*). Co-labeling with pgp9.5 and CGRP revealed that 65-80% of sensory nerves in the retrodiscal tissue of RA, OA, and CH TMJD participants were peptidergic (*Fig. 3B*). Overall, these tissues contained a significantly higher proportion of peptidergic sensory fibers, although the differences between RA, OA, and CH TMJD groups were not statistically significant (*Fig. 3C*).

**Figure 3.**
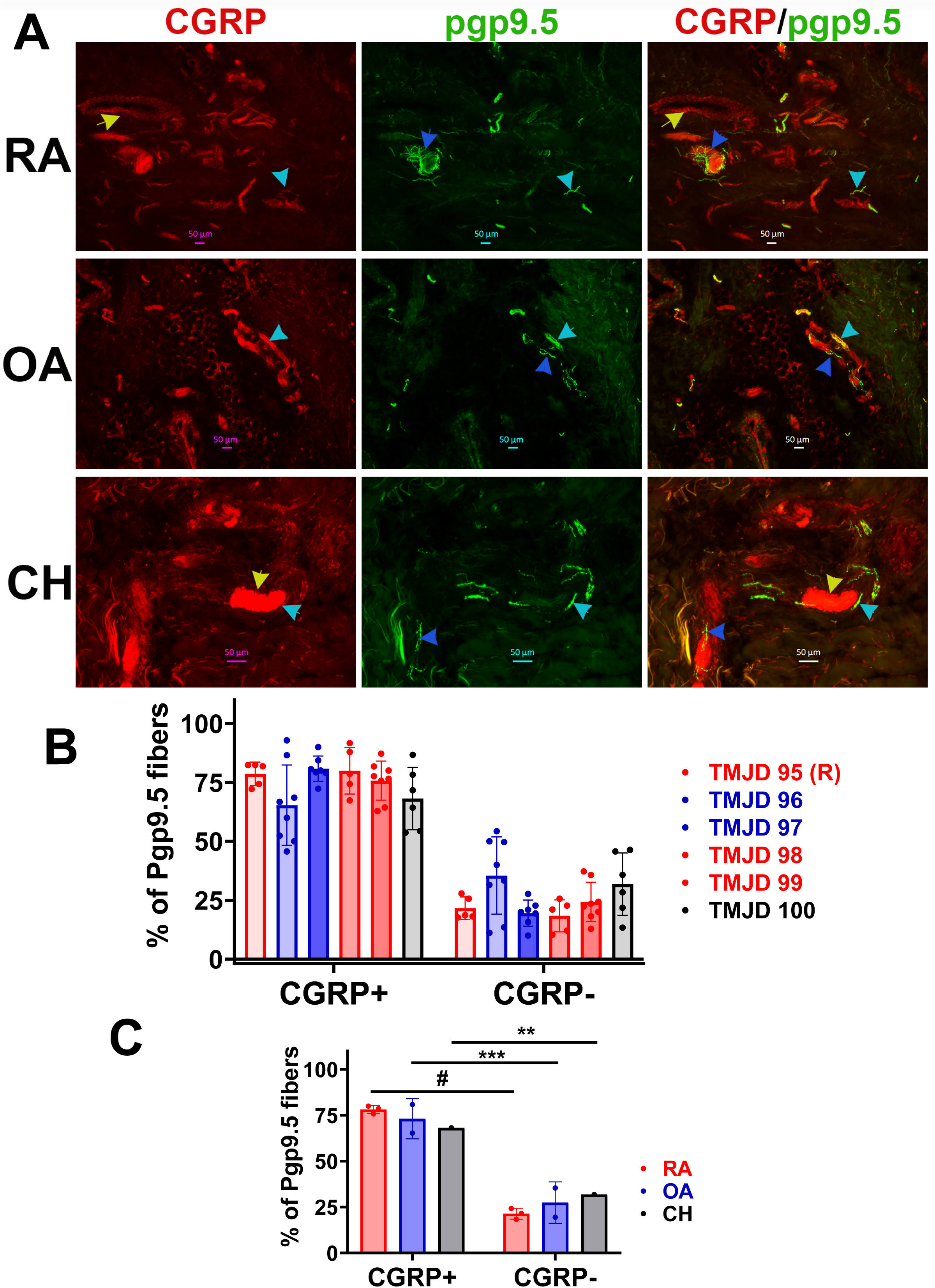
Peptidergic sensory fibers in retrodiscal tissue of TMJD subtypes. **(A)** Representative immunofluorescence micrographs showing CGRP^+^ (peptidergic marker) among pgp9.5^+^ sensory nerve fibers in retrodiscal tissue sections from TMJD participants diagnosed with RA (top row), OA (middle row), and CH (bottom row). Cyan arrows indicate pgp9.5^+^/CGRP^+^ fibers, blue arrows point to pgp9.5^+^/CGRP^−^ fibers, while yellow arrows indicate CGRP^+^ blood vessel. Antibodies and their corresponding fluorescence channels are noted in each image. Scale bars are included in each panel. (**B**) Quantification of pgp9.5^+^/CGRP^+^ and pgp9.5^+^/CGRP^−^ fibers in retrodiscal tissue from individual TMJD participants. Bars for participant #95 show counts from right (R) retrodiscal tissues. RA cases are shown in red, OA in blue, and CH in grey. N as dots on baragraphs represent the number of frames analyzed per tissue. (**C**) Grouped analysis of pgp9.5^+^/CGRP^+^ and pgp9.5^+^/CGRP^−^ fiber percentages across TMJD subtypes. N as dots on bara graphs indicate the number of tissues (or participants) analyzed per group. Statistical significance was determined using two-way ANOVA with Bonferroni’s post hoc test (** p<0.01; *** p<0.001, # p < 0.0001; N=1-3).

Peptidergic fibers can be further classified as myelinated or unmyelinated (Hovhannisyan et al. 2023). Myelinated peptidergic fibers were identified by co-expression of NFH and CGRP, while unmyelinated peptidergic fibers were NFH^−^/CGRP^+^ (*Fig. 4A*). Co-labeling with NFH and CGRP revealed three types of fibers in retrodiscal tissue: NFH^+^/CGRP^+^ (myelinated peptidergic), NFH*-*/CGRP^+^ (unmyelinated peptidergic), and NFH^+^/CGRP^−^ (non-nociceptive myelinated) (Figs. 4A, 4B). Quantification by TMJD subtype showed that NFH^+^/CGRP^−^ fibers - non-nociceptive myelinated nerves - were the least prevalent, comprising only 10–25% of fibers across all TMJD subtypes (*Fig. 4C*). Myelinated and unmyelinated peptidergic fibers were similarly represented, each accounting for 35–50% of fibers in RA, OA, and CH TMJD retrodiscal tissues (*Fig. 4C*). Taken together with the data from Figure 2, these findings indicate that retrodiscal tissues in TMJD contain not only peptidergic nerves but also non-nociceptive (NFH^+^/CGRP^−^) and potentially non-peptidergic unmyelinated fibers. An alternative explanation for the lower observed CGRP labeling could be weak signal intensity below the detection threshold in some NFH^+^ or NFH^−^ nerves.

**Figure 4.**
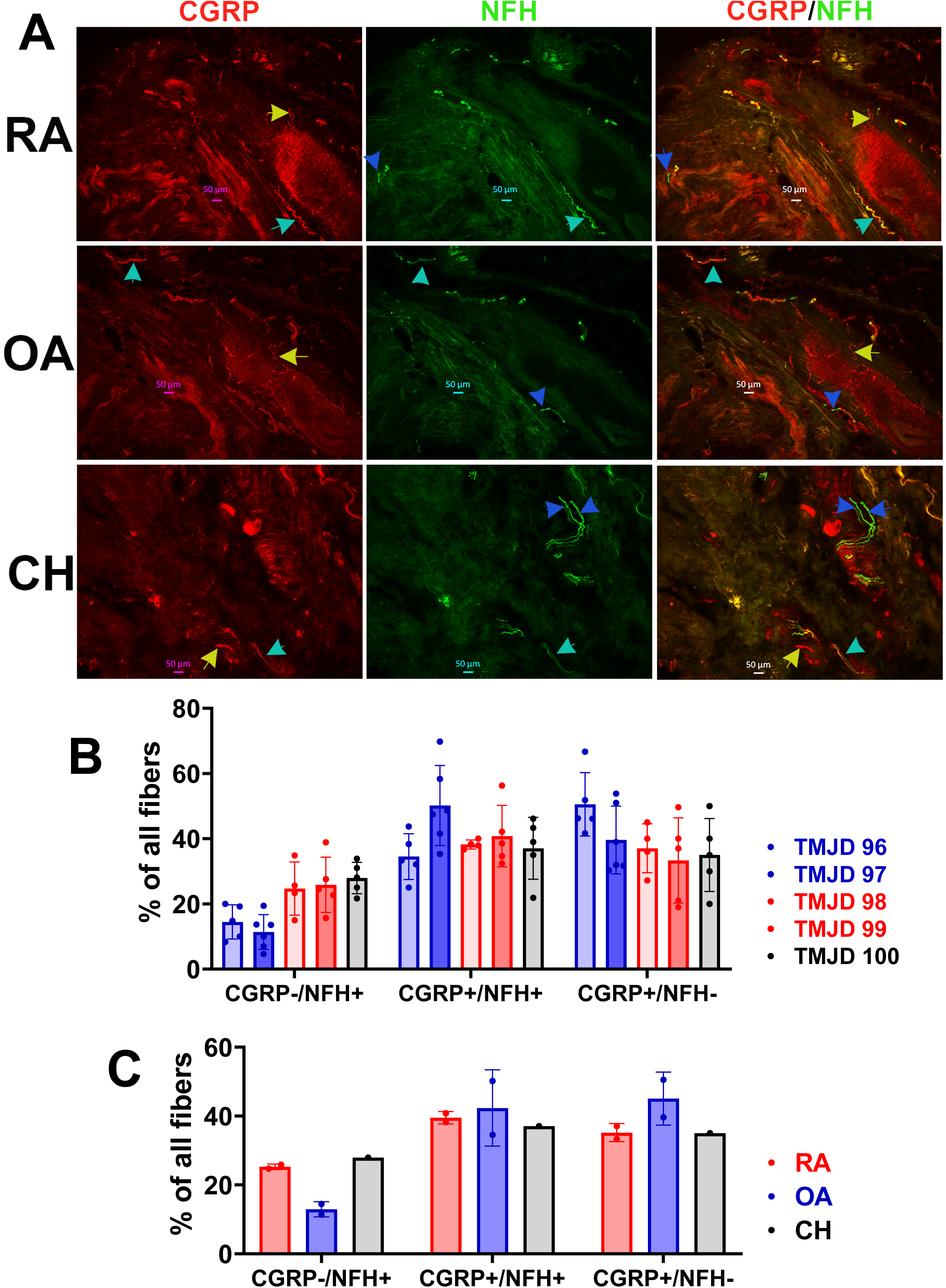
Peptidergic myelinated fibers in retrodiscal tissue of TMJD subtypes. **(A)** Representative immunofluorescence micrographs showing CGRP^+^ among NFH^+^ myelinated nerve fibers in retrodiscal tissue sections from TMJD participants diagnosed with RA (top row), OA (middle row), and CH (bottom row). Cyan arrows indicate NFH^+^/CGRP^+^ fibers, blue arrows point to NFH^+^/CGRP^−^ fibers, while yellow arrows indicate NFH^−^/CGRP^+^ fibers. Antibodies and their corresponding fluorescence channels are noted in each image. Scale bars are included in each panel. (**B**) Quantification of NFH^+^/CGRP^+^, NFH^−^/CGRP^+^ and NFH^+^/CGRP^−^ fibers in retrodiscal tissue from individual TMJD participants. RA cases are shown in red, OA in blue, and CH in grey. N as dots on baragraphs represent the number of frames analyzed per tissue. (**C**) Grouped analysis of NFH^+^/CGRP^+^, NFH^−^/CGRP^+^ and NFH^+^/CGRP^−^ fiber percentages across TMJD subtypes. N as dots on baragraphs indicate the number of tissues (or participants) analyzed per group.

### Innervation of vasculature in retrodiscal tissues from TMJD subtypes

Retrodiscal tissue in the TMJ is known to be highly vascularized (Bouloux et al. 2024a; Bouloux et al. 2024b). To investigate the types of nerves that innervate this vasculature, we analyzed retrodiscal tissues from patients with TMJDs, including RA, OA, and CH. Blood vessels were identified using fluorescently conjugated alpha-smooth muscle actin (α-SMA) antibodies, which label vascular smooth muscle cells and pericytes (Hovhannisyan et al. 2023). Tyrosine hydroxylase (TH) antibodies, typically used to identify sympathetic nerves and a subset of DRG neurons including C-low-threshold mechanoreceptors (C-LTMRs) (Patil et al. 2018; Usoskin et al. 2015), were applied to retrodiscal tissue to mark sympathetic fibers. This choice was based on prior findings that TH rarely labels TG neurons in mice and primates (Hovhannisyan et al. 2023; Lindquist et al. 2021), and that Tafa4 - not TH - is a more appropriate marker for human C-LTMRs in both TG and DRG (Bhuiyan et al. 2024; Tavares-Ferreira et al. 2022). Our data showed that ≥75% of TH^+^ fibers were located in proximity to α-SMA^+^ blood vessels (i.e., within vessel dimensions) across RA, OA, and CH retrodiscal tissues (*Figs. 5A, 5B*). This spatial distribution of TH^+^ fibers was consistent across all TMJD subtypes (*Fig. 5C*).

**Figure 5.**
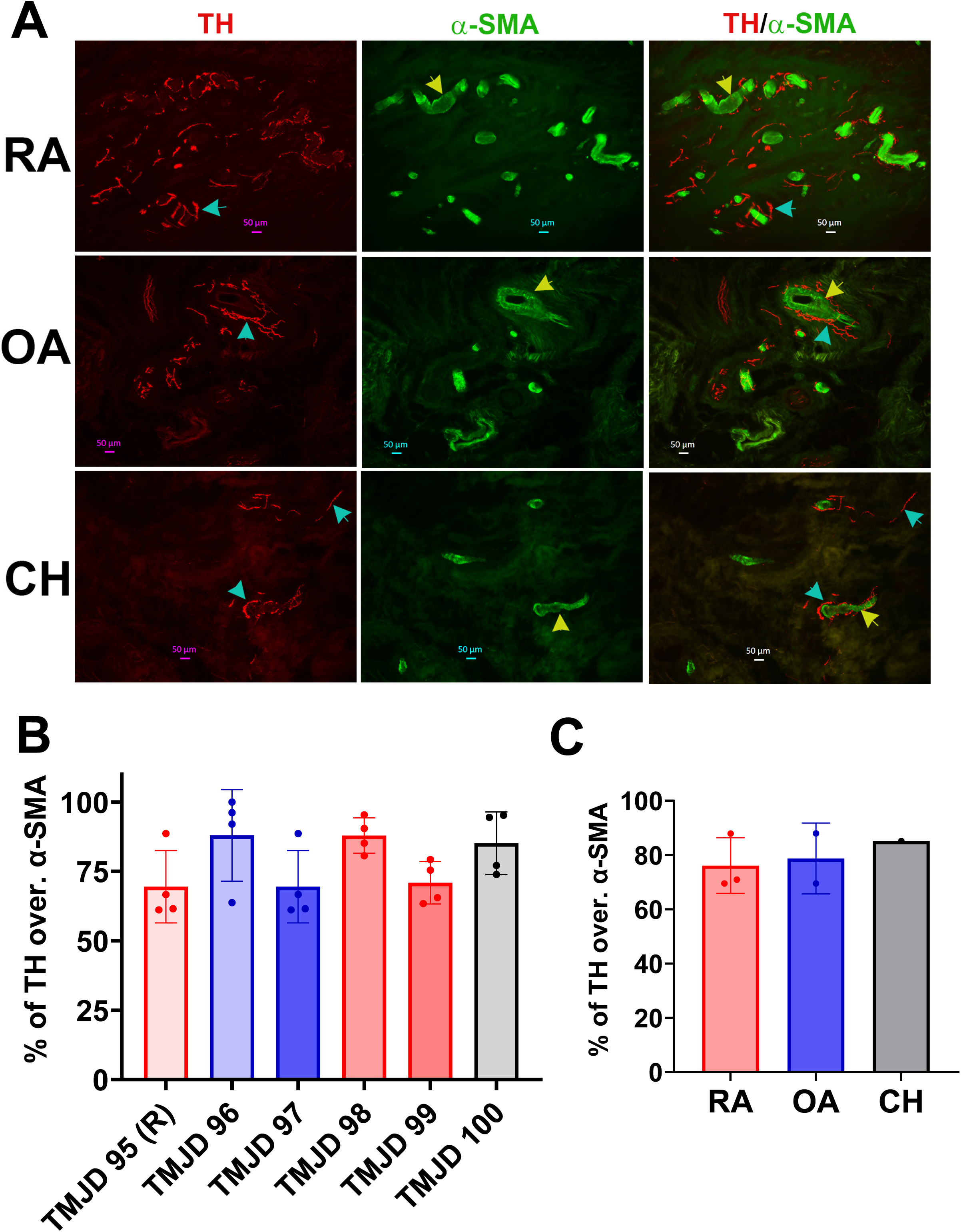
Innervation of blood vessels by sympathetic nerves in retrodiscal tissue of TMJD subtypes. **(A)** Representative immunofluorescence micrographs showing the spatial relationship between tyrosine hydroxylase-positive (TH⁺) sympathetic nerve fibers and alpha-smooth muscle actin-positive (α-SMA⁺) blood vessels in retrodiscal tissue sections from TMJD participants diagnosed with RA (top row), OA (middle row), and CH (bottom row). TH⁺ fibers (cyan arrows) are visualized in close proximity to α-SMA⁺ vasculature (yellow arrows). Fluorescent channels and corresponding antibody labels are indicated in each panel. Scale bars are included. (**B**) Quantification of TH⁺ fibers located near α-SMA⁺ blood vessels in individual retrodiscal tissue samples from TMJD participants. Data for participant #95 include counts from the right (R) joint. RA cases are shown in red, OA in blue, and CH in grey. N denotes the number of frames analyzed per tissue. (**C**) Group-level analysis showing the percentage of TH⁺ fibers in proximity to α-SMA⁺ blood vessels across TMJD subtypes. N represents the number of tissue samples (or participants) per group. Statistical significance was assessed using two-way ANOVA followed by Bonferroni’s post hoc test.

We next examined sensory innervation of vasculature using pgp9.5 as a general sensory neuronal marker. Pgp9.5^+^ sensory fibers were frequently located near α-SMA^+^ vessels, with some fibers running parallel to or wrapping around the vasculature (*Fig. S1A*). Quantitative analysis revealed that 55–75% of pgp9.5^+^ nerves were positioned within the spatial range of α-SMA^+^ vessels (*Fig. S1B*). Although the degree of co-localization varied between individuals, the overall overlap was comparable among RA, OA, and CH groups (*Fig. S1C*).

To determine whether perivascular sensory nerves were myelinated, we conducted immunohistochemistry using NFH antibodies. The majority of NFH^+^ (myelinated) sensory fibers were not found in close proximity to vessels across all TMJD subtypes (*Fig. S2A*), with only 15– 25% located near α-SMA^+^ vessels (*Fig. S2B*). While NFH^+^ fibers sometimes ran alongside vessels, they rarely wrapped around them (*Fig. S2A*). In conclusion, our findings indicate that blood vessels in retrodiscal tissues from RA, OA, and CH TMJD patients are innervated by both sympathetic (TH^+^) and sensory (pgp9.5^+^) fibers. Notably, most sensory fibers associated with vasculature were unmyelinated, a distinction particularly pronounced in RA tissues (*Fig. S3A*).

## Discussion

One of the key events in the development of chronic orofacial pain is the sensitization of sensory neurons (Sessle 2011). Therefore, understanding sensory innervation and neural plasticity in the TMJ is essential for elucidating pain mechanisms in TMJDs. Despite its importance, information on TMJ tissue innervation remains limited, and most existing data come from animal studies or models. In this study, we aimed to define the innervation patterns of the TMJ using tissue samples from human TMJD patients.

TMJDs are broadly categorized into painful and disfunction subtypes (Romero-Reyes and Uyanik 2014; Swift et al. 1998; Tmd (temporomandibular disorders) 2025). RA-related TMJD is a destructive autoimmune condition that causes significant inflammation in TMJ tissues (Savtekin and Sehirli 2018). OA-related TMJD is a degenerative joint disease marked by the breakdown of joints and articular cartilage (Bouloux et al. 2024b; Broussard 2005). Both RA and OA TMJDs are typically associated with intra-articular, bilateral pain and are classified as painful types of TMJD (Bouloux et al. 2024a; Broussard 2005). In contrast, CH is a rare TMJD characterized by facial deformity and functional impairment, and it is classified as a dysfunction-type TMJD (Chouinard et al. 2018). The underlying pathophysiology of RA, OA, and CH differ significantly, suggesting that neural plasticity within the TMJ may also vary across these conditions. Based on this, we selected participants representing these three distinct TMJD types: RA, OA, and CH.

Current understanding suggests that excessive mechanical loading on the TMJ induces initial tissue hypoxia, which activates a cascade of molecular events involving the synovial membrane within the joint capsule (Bouloux et al. 2024a). This hypoxic stress promotes inflammatory cell infiltration and a prolonged inflammatory response, a pivotal process underlying TMJD symptoms such as pain, edema, impaired jaw function, and osteoarthritic changes (Schmidt et al. 2019). These mechanisms highlight the importance of plasticity within TMJ joint capsule. However, the retrodiscal tissue, due to its high vascularity and innervation (Alfaro et al. 2025; Langendoen et al. 1997), may play an equally critical role in initiating the inflammatory cascade. This served as the rationale for focusing our analysis on retrodiscal tissue in our investigation of TMJ innervation.

The primary difference in retrodiscal tissue innervation between painful TMJD subtypes (RA and OA) and the dysfunction subtype (CH) was the density of sensory nerves. Specifically, painful TMJD samples exhibited at least twice the sensory nerve density compared to CH. Despite this quantitative difference, the qualitative distribution of nerve types was largely consistent across RA, OA, and CH cases. In all TMJD subtypes, retrodiscal tissues were predominantly innervated by unmyelinated (NFH^−^) sensory nerves, which accounted for approximately 70% of total sensory fibers. Moreover, most sensory nerves (≈75-80%) were peptidergic, marked by CGRP expression. These peptidergic fibers were nearly evenly split between myelinated (CGRP^+^/NFH^+^) and unmyelinated (CGRP^+^/NFH^−^) subtypes. Based on single-cell RNA-sequencing studies in both human and animal DRG and TG, these nerve populations are consistent with peptidergic C- and Aδ-nociceptors (Bhuiyan et al. 2024). Although C- and Aδ-nociceptors are further subclassified into distinct molecular groups, our study lacked the necessary tools to differentiate these subtypes directly (Bhuiyan et al. 2024). In addition to peptidergic nociceptors, a smaller proportion of retrodiscal nerves were non-peptidergic, including both unmyelinated (CGRP^−^/NFH^−^) and myelinated (CGRP^−^/NFH^+^) fibers. Attempts to label unmyelinated non-peptidergic fibers using established markers such as somatostatin (Sst), MrgprA3, and MrgprD were unsuccessful, even though these markers are known to label such fibers in human skin (Bhuiyan et al. 2024; Tavares-Ferreira et al. 2022). In contrast, labeling with parvalbumin (PV) - a marker of proprioceptors and myelinated non-peptidergic A-low threshold mechanoreceptors (A-LTMRs) - was successful and identified a subset of NFH^+^ fibers in retrodiscal tissue (*Figs. S3C, S3E*). In summary, retrodiscal tissues from TMJD patients harbor a diverse population of sensory nerves. Most are nociceptive and peptidergic in nature, while a smaller subset represents non-nociceptive mechanosensory neurons. This complex innervation pattern underscores the importance of retrodiscal tissue in TMJ sensory signaling and pain mechanisms.

Vasculature within the TMJ plays a critical role in the pathophysiology of TMJDs (Bouloux et al. 2024a; Bouloux et al. 2024b). In light of this, we investigated the innervation of blood vessels in the retrodiscal tissues across TMJD subtypes. Sympathetic innervation of blood vessels plays a critical role in the regulation of vascular tone, blood flow distribution, and blood pressure, all of which are essential for maintaining homeostasis and responding to physiological demands (Cairns 2025; Thomas and Segal 2004). We used TH as a marker for the visualization of sympathetic nerve fibers, as TH is not expressed in human C-LTMRs (Bhuiyan et al. 2024; Tavares-Ferreira et al. 2022). Our findings confirmed robust sympathetic innervation of retrodiscal blood vessels by TH^+^ fibers. In addition to sympathetic fibers, retrodiscal tissue vasculature was also richly innervated by sensory nerves. Notably, most of these sensory fibers were unmyelinated. In contrast, myelinated sensory fibers (NFH^+^) were more frequently observed in regions of the retrodiscal tissue distinct from those occupied by vasculature. However, in some areas, NFH^+^ fibers were found in proximity to TH^+^ sympathetic fibers (*Figs. S3B, S3D*). This distribution pattern contrasts with that observed in the dura mater, where myelinated (NFH^+^) sensory fibers directly innervate blood vessels (Avona et al. 2021).

Despite the careful design and execution of the study, several limitations must be acknowledged. Sex-specific sampling: All tissue samples were obtained from female patients. Previous transcriptomic analyses using bulk and single-cell RNA sequencing have revealed modest, yet distinct, sex-based differences in gene expression within DRG and TG neurons (Bhuiyan et al. 2024; Mecklenburg et al. 2020; Tavares-Ferreira et al. 2022; Yang et al. 2022). Specifically, sex differences were notable in Aβ-rapid adapting (Aβ-RA) and Aδ-low threshold mechanoreceptors (Aδ-LTMRs) under naïve conditions (Barry et al. 2023). However, given the minimal presence of these nerve types in retrodiscal tissues, we do not expect significant sex-based differences in innervation patterns within this context. Manual quantification of nerve fibers: Nerve fiber density was assessed manually from IHC images. This method does not differentiate between closely apposed fibers or separate branches of the same neuron, potentially leading to either overestimation or underestimation of fiber counts. Lack of terminal-specific markers: The study did not include markers that specifically identify neuronal terminals. Therefore, some of the fibers observed and quantified may have been passing through the tissue rather than forming functional terminations within it. Despite these limitations, the central goal of the study - to identify and characterize the predominant and rare nerve fiber types present in retrodiscal tissues across different TMJD subtypes - was achieved. This data provides a foundational map of sensory and sympathetic innervation in these tissues and offers valuable insights into the neurovascular landscape relevant to TMJD pathophysiology.

## Ethical approval and informed consent

Tissues were collected during TMJR from patients enrolled and consented under the “Retrodiscal, Capsular and Discal TMJ Tissue Analysis and Repository” study (IRB #: HSC20220807H) between 2023 and 2025.

## Acknowledgements

We would like to thank Mrs. Priscilla Barba-Escobedo for help in consenting participants. The RE-JOIN consortium consists of Armen Akopian, Kyle Allen, Alejandro Almarza, Benjamin Arenkiel, Maryam Aslam, Basak Ayaz, Yangjin Bae, Bruna Balbino de Paula, Anita Bandrowski, Mario Danilo Boada, Jacqueline Boccanfuso, Jyl Boline, Dawen Cai, Dellina Lane Carpio, Robert Caudle, Racel Cela, Yong Chen, Rui Chen, Brian Constantinescu, Ibdanelo Cortez, Yenisel Cruz-Almeida, M. Franklin Dolwick, Chris Donnelly, Zelong Dou, Joshua Emrick, Malin Ernberg, Danielle Freburg-Hoffmeister, Jeremy Friedman, Spencer Fullam, Janak Gaire, Akash Gandhi, Terese Geraghty, Benjamin Goolsby, Stacey Greene, Nele Haelterman, Zhiguang Huo, Michael Iadarola, Shingo Ishihara, Azeez Ishola, Sudhish Jayachandran, Zixue Jin, Alisa Johnson, Frank Ko, Zhao Lai, Brendan Lee, Yona Levites, Carolina Leynes, Jun Li, Martin Lotz, Lindsey Macpherson, Tristan Maerz, Camilla Majano, Anne-Marie Malfait, Maryann Martone, Simon Mears, Bella Mehta, Emilie Miley, Rachel Miller, Richard Miller, Michael Newton, Alia Obeidat, Soo Oh, Merissa Olmer, Dana Orange, Miguel Otero, Kevin Otto, Folly Patterson, Marlena Pela, Daniel Perez, Sienna Perry, Theodore Price, Hernan Prieto, Russell Ray, Dongjun Ren, Margarete Ribeiro Dasilva, Alexus Roberts, Elizabeth Ronan, Oscar Ruiz, Shad Smith, Mairobys Soccorro Gonzalez, Kaitlin Southern, Joshua Stover, Michael Strinden, Hannah Swahn, Evelyne Tantry, Sue Tappan, Luis Tovias Sanchez, Cristal Villalba Silva, Airam Vivanco-Estella, Robin Vroman, Joost Wagenaar, Lai Wang, Kim Worley, Joshua Wythe, Jiansen Yan, and Julia Younis.

## Author Contributions

J.J.A., A.H.H. and E.E.L.: *methodology, investigation, visualization*. A.H.H., D.E.P. F.J.A. and A.N.A.: *analysis, conceptualization*. D.E.P., F.J.A. and A.N.A.: *research design* J.J.A., RE-JOIN, D.E.P., F.J.A. and A.N.A.– *discussion of results*. A.N.A – *drafted the manuscript*; J.J.A., RE-JOIN; D.E.P., F.J.A. and A.N.A – *discussed and prepared final version of the manuscript*; A.N.A.: *resources, supervision, funding acquisition*. All authors reviewed the manuscript.

## Legends to Supplementary Figures

**Supplementary Figure 1.**
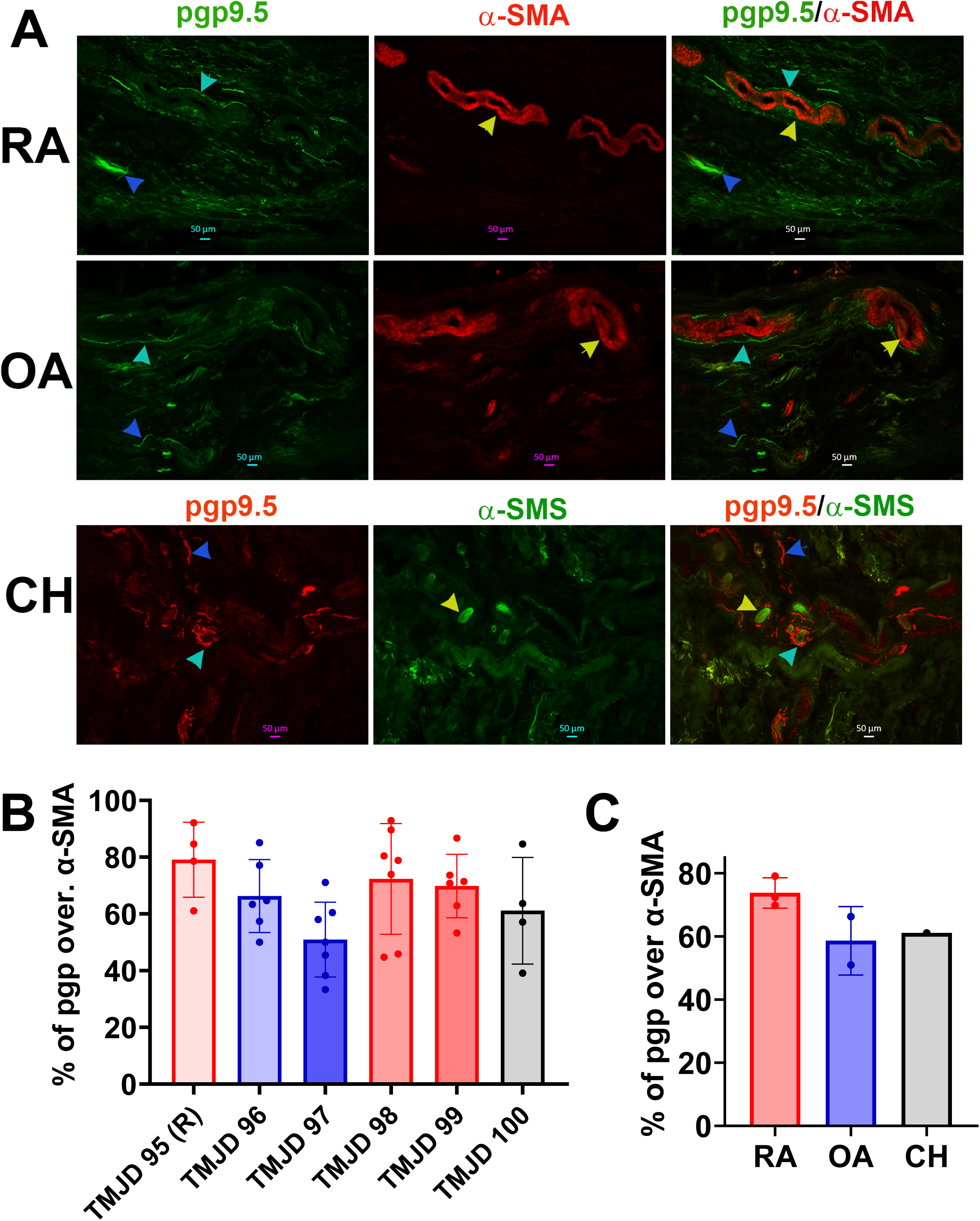
Innervation of blood vessels by sensory nerves in retrodiscal tissue of TMJD subtypes. **(A)** Representative immunofluorescence micrographs showing the spatial relationship between pgp9.5⁺ sensory nerves and α-SMA⁺ blood vessels in retrodiscal tissue sections from TMJD participants diagnosed with RA (top row), OA (middle row), and CH (bottom row). pgp9.5⁺ fibers (cyan arrows) are visualized in close proximity to α-SMA⁺ vasculature (yellow arrows). Fluorescent channels and corresponding antibody labels are indicated in each panel. Scale bars are included. (**B**) Quantification of pgp9.5⁺ fibers located near α-SMA⁺ blood vessels in individual retrodiscal tissue samples from TMJD participants. Data for participant #95 include counts from the right (R) joint. RA cases are shown in red, OA in blue, and CH in grey. N denotes the number of frames analyzed per tissue. (**C**) Group-level analysis showing the percentage of pgp9.5⁺ fibers in proximity to α-SMA⁺ blood vessels across TMJD subtypes. N represents the number of tissue samples (or participants) per group. Statistical significance was assessed using two-way ANOVA followed by Bonferroni’s post hoc test.

**Supplementary Figure 2.**
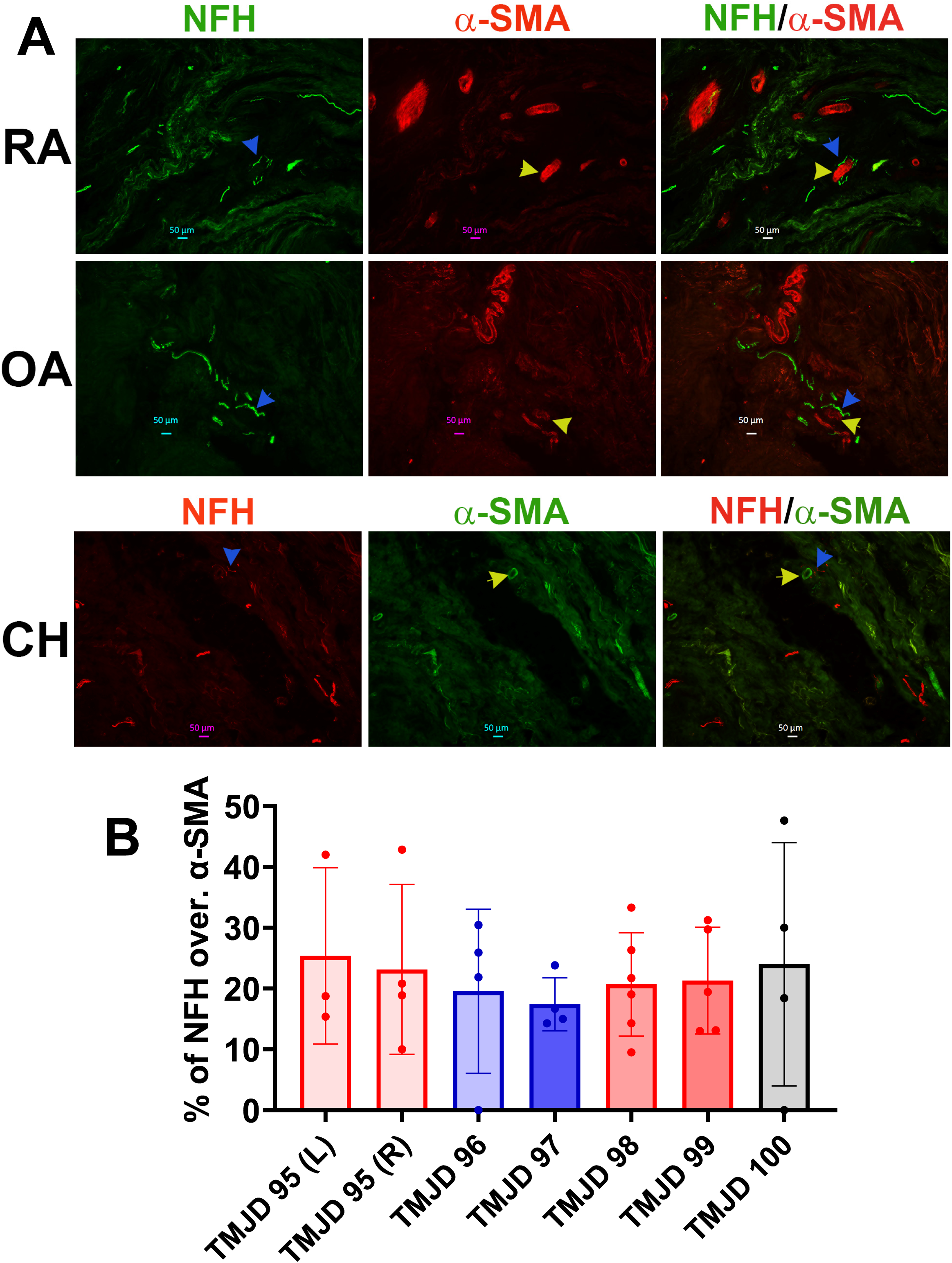
Innervation of blood vessels by myelinated sensory nerves in retrodiscal tissue of TMJD subtypes. **(A)** Representative immunofluorescence micrographs showing the spatial relationship between NFH^+^ myelinated sensory nerve fibers and α-SMA⁺ blood vessels in retrodiscal tissue sections from TMJD participants diagnosed with RA (top row), OA (middle row), and CH (bottom row). NFH⁺ fibers (blue arrows) are visualized in close proximity to α-SMA⁺ vasculature (yellow arrows). Fluorescent channels and corresponding antibody labels are indicated in each panel. Scale bars are included. (**B**) Quantification of NFH⁺ fibers located near α-SMA⁺ blood vessels in individual retrodiscal tissue samples from TMJD participants. Data for participant #95 include counts from both the right (R) and left (L) joints. RA cases are shown in red, OA in blue, and CH in grey. N denotes the number of frames analyzed per tissue.

**Supplementary Figure 3.**
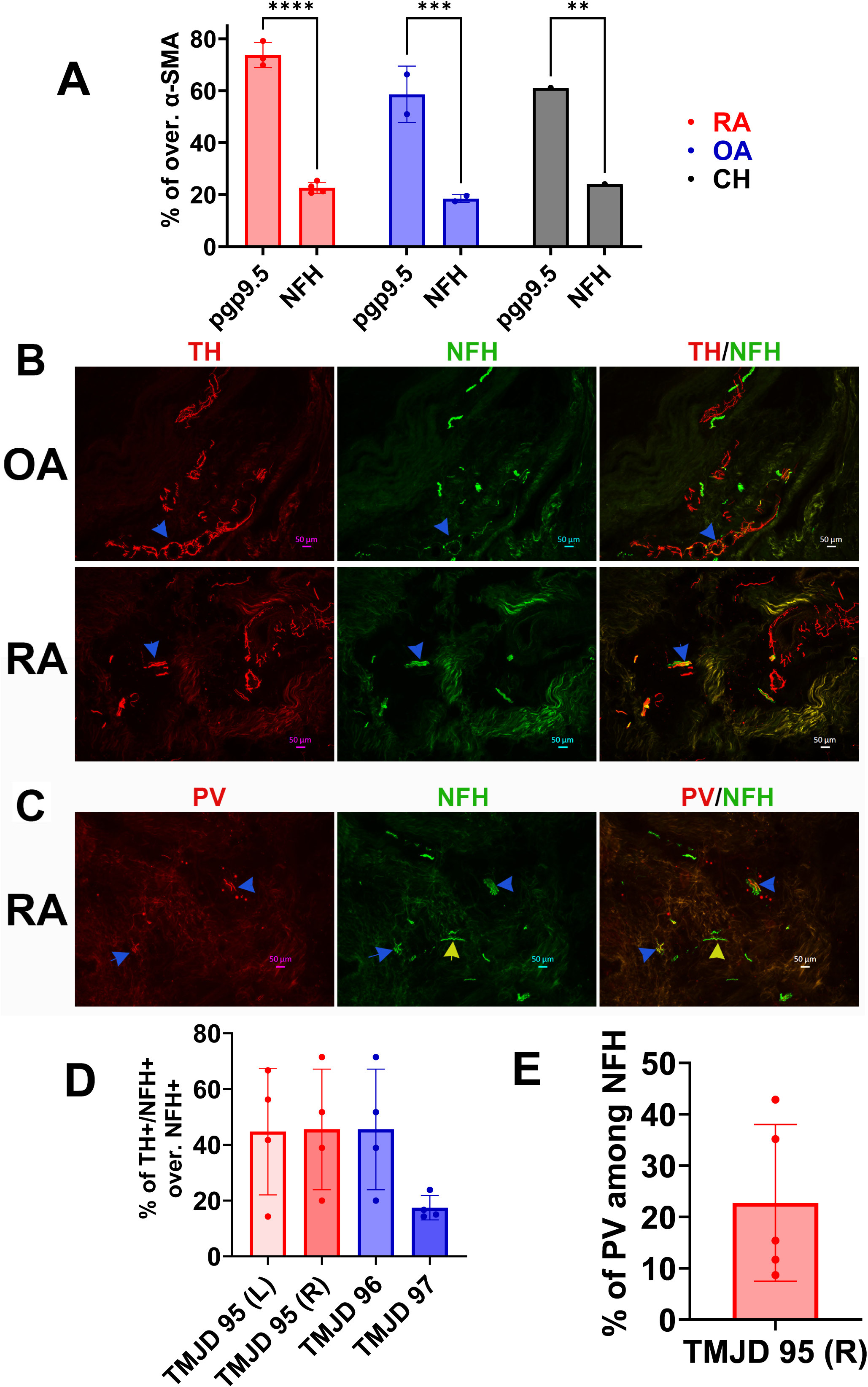
Sensory and sympathetic fibers in retrodiscal tissues. (**A**) Group-level analysis showing the percentage of pgp9.5⁺ compared to NFH⁺ fibers in proximity to α-SMA⁺ blood vessels across TMJD subtypes. N represents the number of tissue samples (or participants) per group. Statistical significance was assessed using two-way ANOVA followed by Bonferroni’s post hoc test (** p<0.01; *** p<0.001; **** p<0.0001). **(B)** Representative immunofluorescence micrographs showing the spatial relationship between TH⁺ sympathetic nerve fibers and NFH⁺ myelinated sensory fibers in retrodiscal tissue sections from TMJD participants diagnosed with OA (top row) and RA (bottom row). Blue arrows point to NFH^+^ fibers overlapping with TH⁺ fibers. Fluorescent channels and corresponding antibody labels are indicated in each panel. Scale bars are included. (**C**) Representative immunofluorescence micrographs showing parvalbumin (PV)^+^ non-nociceptive fibers and NFH^+^ myelinated sensory fibers in retrodiscal tissue sections from a RA TMJD participant. Fluorescent channels and corresponding antibody labels are indicated in each panel. Scale bars are included. (**D**) Quantification of TH⁺/NFH^+^ fibers as percentages of NFH^+^ fibers in individual retrodiscal tissue samples from TMJD participants. Data for participant #95 include counts from both the right (R) and left (L) joints. RA cases are shown in red and OA in blue. N denotes the number of frames analyzed per tissue. (**E**) Quantification of PV⁺/NFH^+^ fibers as percentages of NFH^+^ fibers in the participant #95 right (R) joint.

